# Simultaneous regeneration of skin and bone in full-thickness cranial composite defects

**DOI:** 10.64898/2026.06.16.732662

**Authors:** Mirae Kim, Yi Zhu, Shivakalyani Adepu, Caralyn P. Collins, Maria Mendez-Santos, Cheng Sun, Tong-Chuan He, Russell R. Reid, Guillermo A. Ameer

## Abstract

Traumatic cranial defects often involve concurrent loss of soft and hard tissues and can progress to chronic defects due to delayed healing associated with infection or other co-morbidities. Despite autologous reconstruction remaining the clinical standard, it requires staged procedures using heterogeneous tissues, increasing operative time, costs, and surgical risks. Moreover, current tissue engineering approaches focus on single tissues or acute tissue defect models, limiting their clinical applications. Herein, we describe an acellular, material-driven 3D-printed composite scaffold designed to regenerate both bone and skin within composite cranial defects. The scaffold integrates controlled copper ion release from both organic and inorganic components with 3D-printed citrate polymer and citrate polymer–ceramic composites. Integrated thermoresponsive citrate-based hydrogels further enable spatially defined dermoconductive and osteoconductive properties, supporting a one-step surgical approach. At 12 weeks post-implantation, our scaffold enhanced keratinocyte organization, collagen deposition, and defect coverage with mature bone, achieving histological outcomes comparable to autografts. Furthermore, the system suppressed bacterial burden. Thus, this acellular platform represents a clinically promising synchronized strategy to address the complex demands of traumatic craniofacial composite defects.

## 1. Introduction

High-energy penetrating trauma often results from ballistic injuries in military or civilian settings and involves damage across multiple tissues with complex three-dimensional structures.^[1]^ A substantial proportion of fatal combat injuries are associated with high-energy craniomaxillofacial (CMF) trauma, which accounts for approximately 40% of high-level (Level III or higher) patient evacuations in combat environments.^[2]^ More than 75% of these cases involve open facial defects with extensive destruction of the skull, scalp, and surrounding soft tissues, with reported mortality rates reaching up to 78%, primarily due to infection, cerebrospinal fluid leakage, and severe hemorrhage.^[3]^

While autologous tissue, in the form of grafts and flaps, remains the clinical gold standard, their application in CMF reconstruction is often limited for large or complex defects due to the need to replicate intricate craniofacial anatomy, limited tissue availability, donor-site morbidity, and high rates of bone graft resorption.^[4, 5]^ Furthermore, although single-tissue defects can often be managed using standardized treatment, complex defects involving multiple tissue types typically require multi-stage reconstructive surgeries or complex free tissue transfers.^[6, 7]^ These multistage procedures substantially increase operative time and cost, require advanced surgical expertise, and further elevate the risk of complications and adverse outcomes.^[6, 8]^

Regenerative engineering has been considered as a promising alternative.^[9]^ An ideal system supports mechanically competent bone regeneration, enables rapid and stable soft-tissue coverage, and facilitates vascular reconstruction across complex tissue interfaces in a spatiotemporally coordinated manner.^[10, 11]^ However, most existing scaffold-based approaches are designed to address only one aspect of this multifaceted repair process or a single tissue type. In addition, there is a lack of studies that adequately recapitulate clinically relevant defect conditions, making it difficult to evaluate therapeutic efficacy.^[12]^ For example, in combat settings, cranial reconstruction is often delayed, up to three months, resulting in a chronic defect environment characterized by impaired vascularity, hypoxia, persistent inflammation, and fibrosis.^[7, 13]^ These conditions differ fundamentally from acute defect models and can lead to different regenerative outcomes.^[14]^ This limitation is further exacerbated in systems incorporating exogenous cells or growth factors, which often introduce additional regulatory complexity, and potential immune responses, thereby complicating timely and predictable reconstruction. Importantly, the inclusion of such exogenous components also complicates clinical translation due to increased manufacturing challenges, regulatory burden, and cost.^[15]^

Bioactive glass (BG) has been recognized as a clinically validated material for bone regeneration since its FDA approval in 1985.^[16, 17]^ Copper, an essential trace element, supports immunomodulation, angiogenesis, extracellular matrix remodeling, and antimicrobial defense, making it a versatile bioactive cue for tissue repair.^[18, 19]^ Hydroxyapatite (HA), a naturally occurring calcium apatite mineral, is widely used for its biocompatibility, osteoconductivity, and similarity to native bone mineral. Citrate-based elastomers enhance bone regeneration by promoting mesenchymal stem cell osteogenesis through citrate-mediated metabolic regulation.^[20]^ FDA-clearance of citrate-based material/HA composites for musculoskeletal applications further supports the clinical translatability of this materials platform.

Herein, we introduce a citrate-based acellular integrated platform for complex cranial defects, based on a materials strategy offering clinical and regulatory advantages. In our previous work, we demonstrated that a peptide-conjugated (A5G81–PPCN) or copper ion-eluting (CMOF-PPCN) hydrogel enhanced chronic wound healing^[21-23]^, and a personalized mineral scaffold (mPOC–60HA) promoted bone regeneration in critical-sized bone defect^[5]^. Building upon our previous work, we engineered a single-piece, 3D-printed multizonal scaffold–hydrogel composite designed for concurrent hard and soft tissue regeneration (**Figure 1**). The printed scaffold fabricated from citrate-based elastomers provides structural support across bone and soft-tissue regions, while a citrate-based thermoresponsive hydrogel (PPCN) enables personalized conformity to irregular defect geometries. Copper was selectively incorporated in a tissue-specific manner. Copper-doped bioactive glass (BGCu) was integrated into the bone-directed hydrogel, while a copper metal–organic framework (CMOF) was incorporated into the skin-directed A5G81-PPCN hydrogel. This single-stage construct was designed to spatially coordinate regenerative cues within a unified platform, enabling synchronized healing of heterogeneous tissues. Our results show that, in a clinically relevant chronic defect model^[24]^, the copper ion-eluting system achieves regenerative outcomes comparable to autografts with only material-driven strategy. These findings highlight our unified, acellular system as a clinically promising platform for rapid stabilization and coordinated multi-tissue regeneration in traumatic craniofacial composite defects.

**Figure 1.**
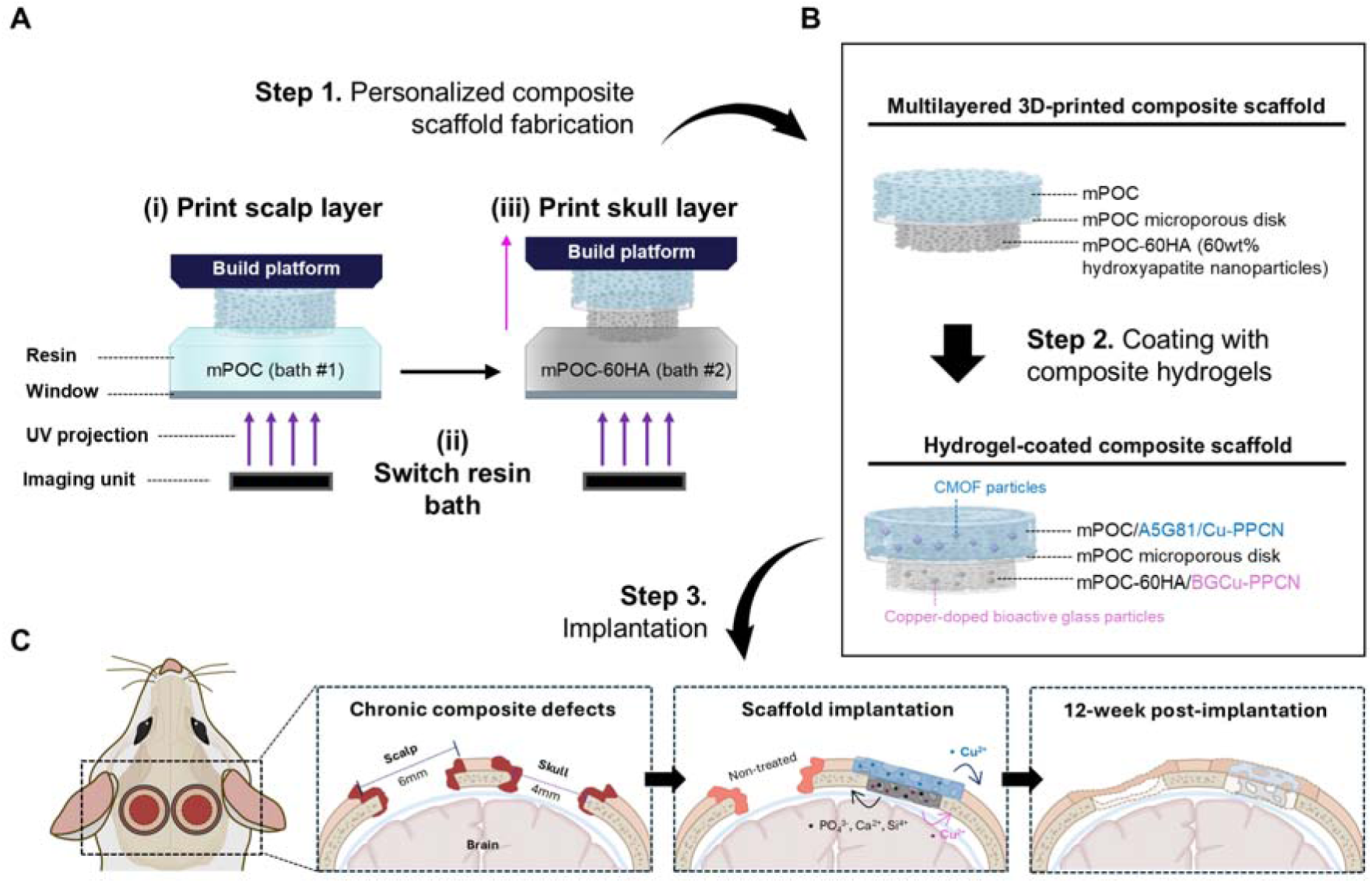
Schematic illustration of the fabrication and implantation process of the multizonal composite scaffold. **(A)** Multizonal composite scaffold fabrication by sequential printing of skin and bone regions using resin bath switching, **(B)** followed by thermoresponsive hydrogel (PPCN) coating, and **(C)** implantation into a chronic composite defect in a rat model.

## 2. Results

### 2.1. A multizonal composite scaffold integrates soft and hard tissue regions within a single construct

A single-piece multizonal composite scaffold was fabricated using a micro-continuous liquid interface production (µCLIP) 3D printing system^[25]^ (**Figure 1A**). The construct consisted of a soft, polymeric scalp region and a mineralized skull region integrated within a single continuous structure. The scalp zone was composed of mPOC (**Figure S1**), providing flexible structural support, whereas the skull zone was formed from mPOC reinforced with 60 wt% HA (mPOC-60HA) to approximate the mineral density of native bone. Between the two regions, a thin microporous mPOC interfacial layer was incorporated to enable gradual transition across the composite tissues.

Following printing, the entire construct was uniformly coated with composite PPCN hydrogels at 37 °C (**Figure 1B and Figure S2, S3).** This approach yielded a single implant that combined spatially distinct mechanical and biochemical cues within one integrated scaffold. The resulting hydrogel-coated multizonal scaffolds maintained structural integrity and were readily handled and implanted into bilateral composite cranial defects (**Figure 1C**).

### 2.2. Composite PPCN hydrogels enable zone-specific coating and sustained copper ion release

Each scaffold zone was designed with hexagonal unit structure, and the 3D-printed multizonal composite scaffold system exhibited seamless integration between different materials across adjacent zones (**Figure 2A**).

**Figure 2.**
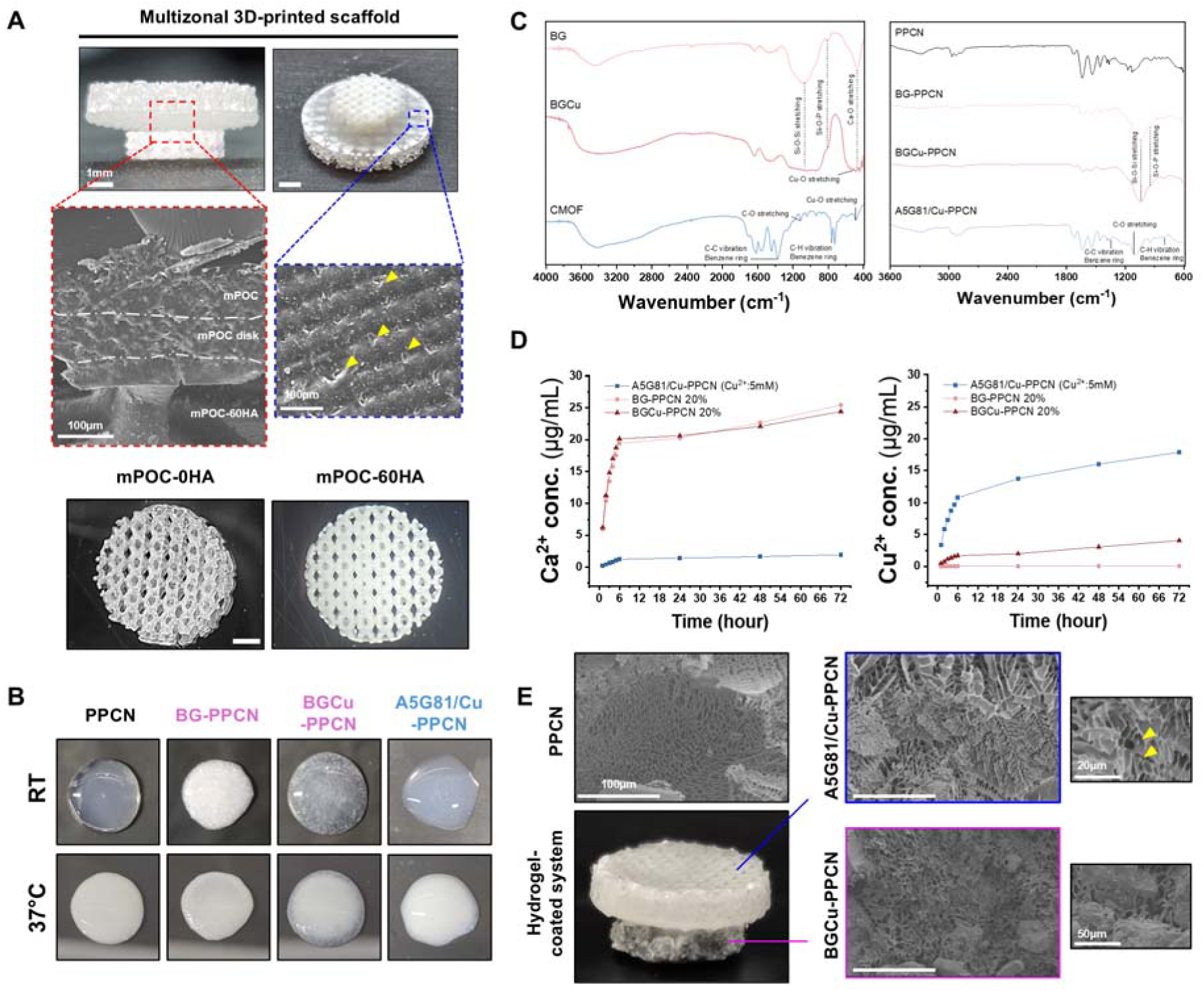
Characterization of the composite PPCN hydrogels and fabrication of hydrogel-coated 3D-printed scaffolds. **(A)** Photographs of 3D-printed scaffolds without HA (mPOC-0HA) and with 60 wt% HA (mPOC-60HA), along with SEM images showing the morphological structure of each printed zone. **(B)** Thermoresponsive behavior of each composite PPCN hydrogel at room temperature and 37□°C. **(C)** FT-IR spectra of the composite hydrogels and the individual composite particles. **(D)** Calcium ion and copper ion release profiles from composite PPCN hydrogels over time. **(E)** Photograph of the hydrogel-coated composite scaffold and SEM images showing the internal morphology of the composite hydrogel.

The compositions of BG, BGCu and CMOF was confirmed via energy-dispersive X-ray spectroscopy, and the crystallographic structure of CMOF was verified using X-ray diffraction analysis (**Figure S4, S5**). BG and BGCu exhibited comparable elemental compositions in Si (24.17 and 23.30 wt%), Ca (19.52 and 20.22 wt%), and P (3.00 and 3.05 wt%), while copper was doped in BGCu at approximately 1.03 wt%. In the CMOF particles, copper accounted for approximately 22.42 wt% (based on total C, O, and Cu content).

All composite PPCN hydrogels maintained the thermoresponsive behavior characteristic of PPCN (**Figure 2B**), exhibiting a gelation temperature in the range of 30.5–32.23 °C (**Figure S6**). The elemental composition (Si, Ca, P, and Cu) was comparable between BG- and BGCu-PPCN hydrogels (**Figure S7**), facilitating accurate assessment of copper-specific effects in subsequent experiments. The composite PPCN hydrogels exhibited characteristic FT-IR peaks corresponding to each component (**Figure 2C**). Si–O–Si and Si–O–P bonding vibrations appeared at approximately 1033.87 cm□¹ and 939.35 cm□¹, respectively. In addition, C–C and C–H vibrational peaks derived from the CMOF particles were detected near 1348.27 cm□¹ and 802. 40 cm□¹.

The ion release profiles of the composite PPCN hydrogels were evaluated using ICP–MS at an elevated concentration (**Figure 2D**). Calcium release from BG- and BGCu-PPCN hydrogels followed a similar trend, reaching approximately 4.04 µg/mL after 72 h, thereby allowing direct comparison of the copper-dependent effects. In A5G81/Cu-PPCN hydrogels, copper release reached 17.9 µg/mL at 72 h. These results indicate that PPCN hydrogels support sustained release of both copper and calcium ions, thereby minimizing the risk of rapid local ion accumulation.^[22]^

The thermoresponsive properties of PPCN enabled efficient handling and uniform coating of the composite hydrogels according to the intended tissue region (**Figure 2E**). SEM imaging further revealed that each composite hydrogel exhibited a highly porous architecture, which is expected to facilitate tissue ingrowth.

### 2.3. Released Cu²□ modulates cell behavior to enhance osteogenesis and wound closure

Based on the observed cell viability in hydrogel-coated composite scaffold, a 10 wt% concentration of BG or BGCu in the PPCN hydrogel was identified as optimal and was therefore selected for subsequent experiments (**Figure S8**). Likewise, a Cu^2+^ concentration of 1 mM within the A5G81/Cu-PPCN hydrogel was chosen, as our previous study demonstrated that controlled release of Cu²□ from this formulation supported cell viability and promoted wound healing.^[22]^

To assess the effects of BG- and BGCu-PPCN hydrogels, osteogenic marker expression was examined at days 7 and 14 in human mesenchymal stem cells (hMSCs) cultured on mPOC-60HA scaffolds with or without hydrogel coatings, using immunofluorescence staining (**Figure 3A**). At day 7, hydrogel-coated scaffolds exhibited significantly higher expression of OPN, OCN, and RUNX2 compared to uncoated scaffolds (**Figure 3B**). In particular, BGCu-PPCN–coated scaffolds showed markedly higher levels of OPN (2.60 ± 0.11, p < 0.001) and OCN (3.42 ± 0.22, p < 0.0001) compared to BG-PPCN–coated scaffolds. At day 14, OPN and OCN expression elevated in both uncoated and BG-PPCN–coated scaffolds, with higher levels observed in the BG-PPCN group. In contrast, in the BGCu-PPCN group, OPN expression (2.47 ± 0.13) showed minimal change, while OCN and RUNX2 expression decreased substantially. These findings indicate that both BG- and BGCu-PPCN hydrogels promoted osteogenic differentiation of hMSCs. Moreover, released copper ions accelerated progression toward a more mature osteogenic stage, likely promoting extracellular matrix deposition rather than prolonged expression of early-osteogenic markers.^[26, 27]^

**Figure 3.**
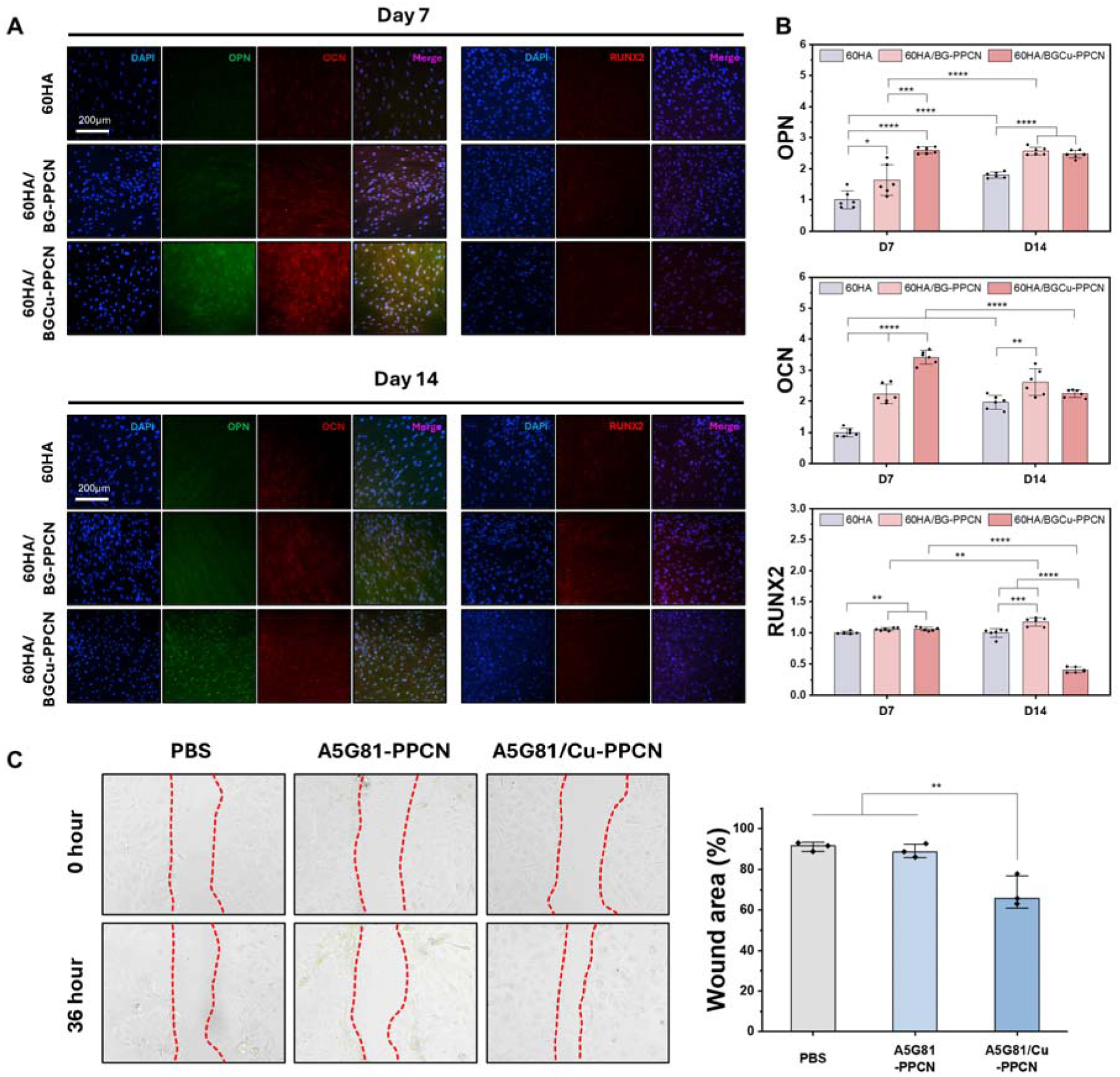
*In vitro* evaluation of osteogenic differentiation and wound healing responses to copper-eluting hydrogels. **(A)** Representative immunofluorescence images of osteogenic markers at days 7 and 14. **(B)** Quantification of marker expression based on mean gray value at days 7 and 14. **(C)** Digital images showing HEKa cell migration after treatment with PBS, A5G81-PPCN, and A5G81/Cu-PPCN hydrogels, along with quantification of wound closure area after 36 h.

Wound healing capacity was evaluated using human epidermal keratinocytes (HEKa) in a scratch assay (**Figure 3C**). After 36 hours, no significant difference was observed in the A5G81-PPCN hydrogel group compared with the PBS-treated group. Notably, treatment with A5G81/Cu-PPCN hydrogels resulted in 31.02 ± 6.48% (p < 0.01) wound closure, indicating that copper released from CMOF particles accelerated the wound healing activity.

### 2.4. Cu^2+^-eluting PPCN hydrogels enhance vascular maturation and antibacterial activity

Tubule formation was assessed using human umbilical vein endothelial cells (HUVECs). After 24 hours of incubation with each composite PPCN hydrogel, the numbers of tubule nodes, junctions, and meshes were quantified (**Figure 4A**). Although no significant differences were observed in tubule nodes between non-copper hydrogels (BG-PPCN and A5G81-PPCN) and copper ion-eluting hydrogels (BGCu-PPCN and A5G81/Cu-PPCN), released copper ions significantly increased the number of junctions and meshes (p < 0.05). These findings suggest that copper ions primarily promote vascular maturation by enhancing network complexity and stabilizing forming tubules, rather than by increasing tubule branching.^[26]^

**Figure 4.**
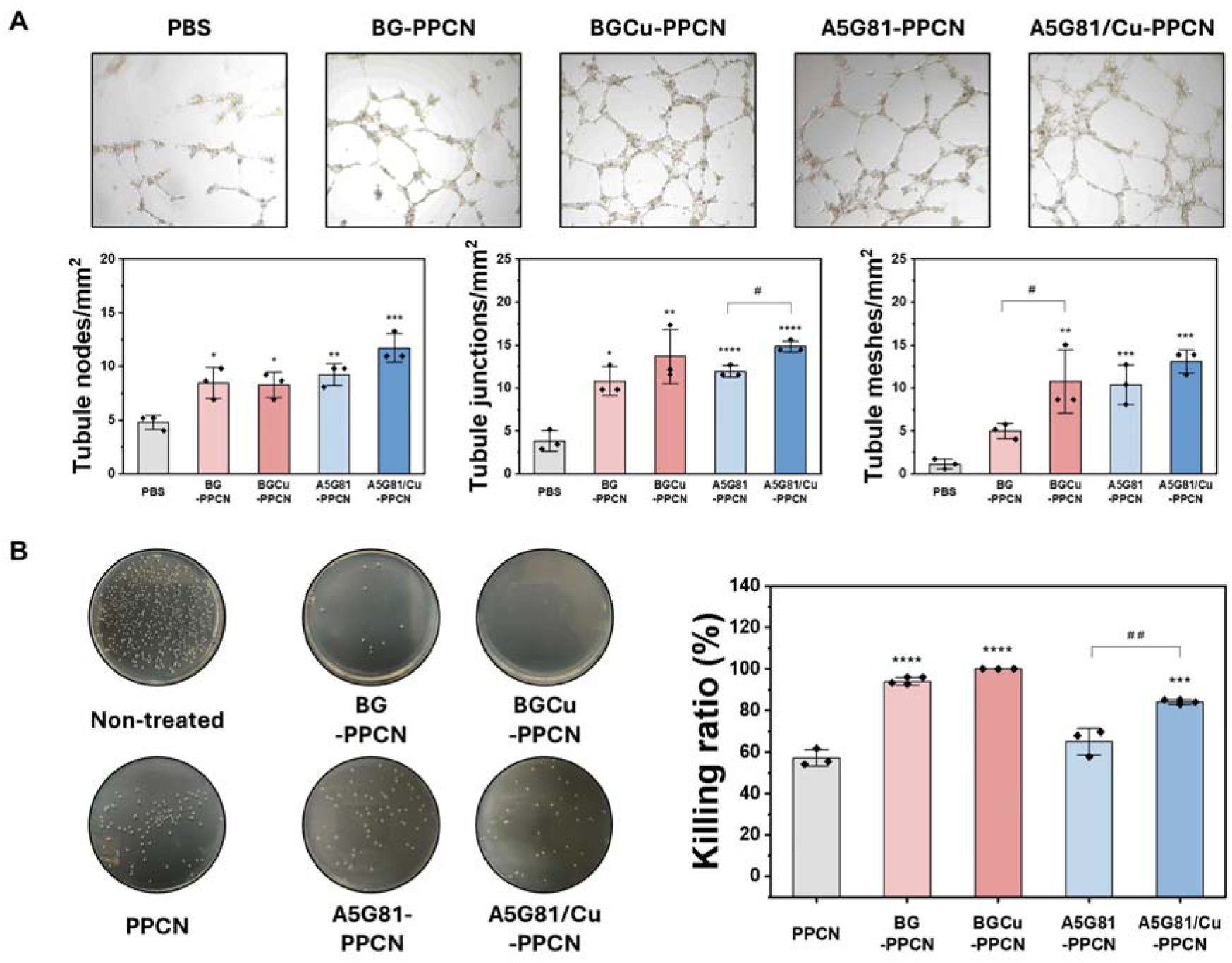
*In vitro* assessment of angiogenic and antibacterial properties of copper-eluting hydrogels. **(A)** Representative images of HUVEC tube formation after 24 hour incubation with each hydrogel, with quantification of tube nodes, junctions, and meshes per unit area. **(B)** Images of S. aureus colonies on agar plates after 24 hour incubation with each hydrogel. * indicates statistical significance vs. PBS or PPCN; # indicates comparisons within the same group.

To evaluate the antibacterial efficacy of the composite hydrogels, we assessed their activity against *Staphylococcus aureus* (*S. aureus*), the most common bacterial species associated with chronic wounds and post-traumatic cranial infections^[28, 29]^ (**Figure 4B**). PPCN hydrogel alone demonstrated a killing ratio of 57.18 ± 3.99%, likely due to its ability to inhibit bacterial growth through metal ion chelation (e.g., calcium and magnesium ions) and subsequent disruption of the bacterial cell membrane.^[30]^ The A5G81-PPCN hydrogel (65.19 ± 6.45%) did not show a significant improvement compared to PPCN alone (p = 0.141). In contrast, A5G81/Cu-PPCN exhibited a markedly enhanced antibacterial effect, with a killing ratio of 84.03 ± 1.12% (p < 0.01 vs. A5G81-PPCN). Similarly, hydrogels containing bioactive glass (BG-PPCN and BGCu-PPCN) also showed increased bacterial suppression, attributed to ion release from the bioactive glass particles.^[31]^ However, no significant difference was observed between the two formulations.

Collectively, copper ion-eluting PPCN hydrogels not only enhanced tubule formation and vascular maturation but also significantly improved antibacterial activity.

### 2.5. Cu^2+^-eluting scaffolds accelerate calvarial bone and scalp soft tissue regeneration

To evaluate composite defect regeneration involving both scalp and skull, bilateral chronic defects were created in rats based on our previously established model.^[24]^ For each animal, scaffold treatments were implanted on one side, while the contralateral side served as a control (**Figure 5A**) allowing comparison of tissue regeneration under identical biological and environmental conditions. The positive control consisted of autologous tissue. Specifically, the skull defect was immediately covered with the craniotomy bone piece and the scalp defect was closed with the rotational flap. The negative control was left untreated. To evaluate the effects of Cu²□ release from the scaffold, the w/o Cu² group consisted of scaffolds coated with non-copper hydrogels, while the w/ Cu²□ group consisted of scaffolds coated with copper ion-eluting hydrogels.

**Figure 5.**
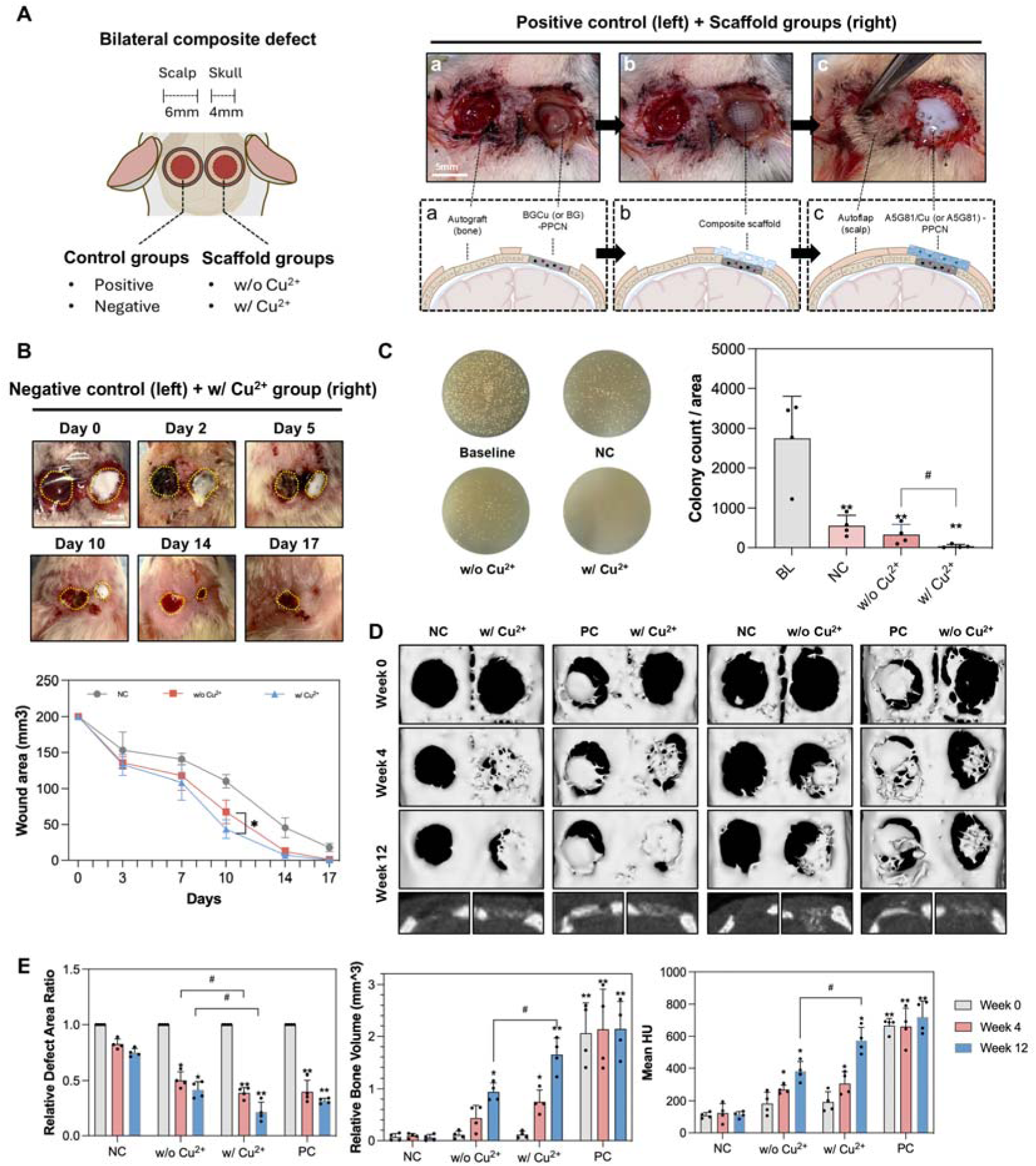
Composite defect regeneration using the hydrogel-coated multizonal scaffold system in a critical-sized chronic cranial defect model. **(A)** Schematic illustration of the experimental setup and intraoperative images of scaffold implantation. a) Autologous bone graft implanted on the left side, with application of the bone-region composite hydrogel on the right side. b) Placement of the multizonal composite scaffold on the right side. c) Coverage of the left side with a local pedicled skin flap (autoflap) and application of the skin-region composite hydrogel on the right side. **(B)** Representative images of wound healing progression over time in the negative control and copper-eluting scaffold groups, observed until complete closure. **(C)** Bacterial colonization in the defect area on day 5 post-treatment for each group. **(D)** Representative micro-CT images at 0, 4, and 12 weeks post-treatment. **(E)** Quantitative analysis of relative defect area, regenerated bone volume, and mean Hounsfield units (HU) over time for each treatment group. NC: negative control (untreated); PC: positive control (autologous tissues); w/o Cu^2+^: scaffolds coated with non-copper hydrogels; w/ Cu^2+^: scaffolds coated with copper-eluting hydrogels. * indicates statistical significance vs. NC; # indicates comparisons within the same group.

Scalp wound closure was monitored over time and compared with the negative control group (**Figure 5B**). No visual or quantitative differences were observed between treatment groups until day 3. However, from day 10, the w/ Cu^2+^ groups exhibited significantly accelerated wound closure, achieving approximately 70% closure compared to 62.5% in w/o Cu^2+^ groups (p = 0.039) (**Figure S9**). By day 14, the w/ Cu^2+^ groups achieved approximately 100% wound closure, whereas the untreated control remained at ∼70% and reached complete closure only by day 21 (data not shown).

On day 5, when defects were still partially open and intergroup differences were minimal (p= 0.376; quantified data not shown), wound swabs were cultured on agar plates to assess bacteria-suppressing effect during the healing process (**Figure 5C**). Compared to the baseline (collected on day 0), all treatment groups exhibited reduced bacterial growth. Notably, compared with the negative group, the w/ Cu^2+^ group exhibited a substantially reduction in bacterial burden (killing ratio 98.7%), whereas the w/o Cu^2+^ group showed a more modest effect (killing ratio 87.5%).

New bone formation was evaluated using micro-CT at week 0, week 4 and week 12. Three-dimensional micro-CT reconstruction (**Figure 5D**) revealed that at week 0, bilateral cranial defects were of comparable size across groups, and the positive group showed re-implanted autologous bone. By week 4, the negative group exhibited modest reduction in defect size, both w/ Cu^2+^ and w/o Cu^2+^ groups showed partial new bone formation, while the w/ Cu^2+^ group displaying a markedly larger bone volume. At week 12, the negative group defect size remained largely unchanged compared to week 4, suggesting that the peak healing of critical-sized defects occurs within the first 4 weeks.

The relative defect area ratio, bone volume, and mean Hounsfield unit (HU) intensity were calculated based on micro-CT datasets (**Figure 5E**). Statistical analysis indicated that the positive group exhibited the most effective cranial defect repair. Both scaffold groups showed significant improvements in defect coverage, bone volume, and bone mineral density compared to the negative group. Notably, total bone volume differed significantly between scaffold groups at week 12 (p= 0.036), and the w/ Cu^2+^ group demonstrated greater defect coverage (∼ 1.4-fold) and higher mean HU values (∼1.5-fold) than the w/o Cu^2+^ group at 12 weeks. These results suggest that copper enhanced structural organization, integration with native bone, and formation of more mineral-dense tissue.

### 2.6. Sustained copper-ion release enables mature composite tissue regeneration that is comparable to autografts

At 12 weeks post-implantation, tissues were collected and histologically evaluated by H&E and Masson’s trichrome staining (**Figure 6A**). In the scalp region, all treated groups demonstrated comparable skin regeneration, whereas the negative control showed lower collagen deposition density. In particular, in the skull region, the w/ Cu^2+^ treated groups exhibited substantially more mature bone formation along the scaffold, demonstrating improved osseointegration. In contrast, the w/o Cu^2+^ scaffolds induced limited osteoid tissue formation surrounding the scaffold structure.

**Figure 6.**
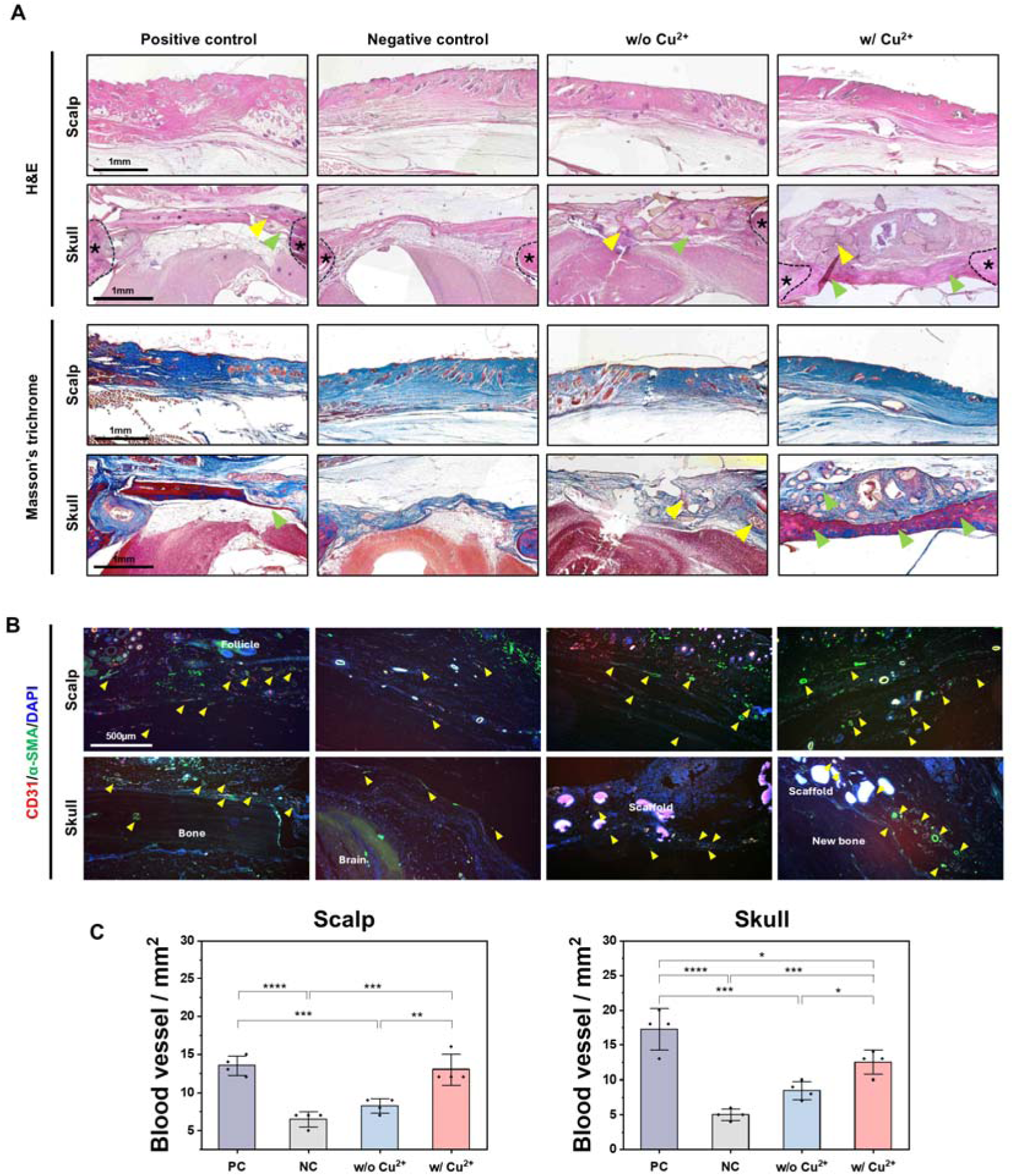
Histological evaluation of the treatment groups in the composite cranial defect model. **(A)** Representative H&E and Masson’s trichrome staining of scalp and skull defect regions at 12 weeks post-implantation. Yellow arrow: osteoid; green arrow: newly formed bone; asterisk: native bone; black dash line: defect site. **(B)** Assessment of angiogenesis in each group by CD31 and α-SMA immunostaining in the scalp and skull regions. **(C)** Quantitative assessment of blood vessel formation in the scalp and skull regions. Yellow arrow: blood vessels.

Blood vessel formation within the newly regenerated tissue was quantified using CD31 and α-SMA co-stained sections (**Figure 6B**). In the scalp region, the w/ Cu^2+^ group exhibited markedly enhanced vascularization (13.0 ± 2.3/mm2) compared with both the negative control (6.5 ± 1.0/mm2) and the w/o Cu^2+^ treatment group (8.25 ± 0.9/mm2) (**Figure 6C).** In contrast, the w/o Cu^2+^ group only showed a significant increase compared with the negative control in the skull region.

The regenerated tissue quality in the scalp region was assessed using collagen type I (dermal marker) and keratin 10 (epidermal marker) immunostaining (**Figure 7A**). The w/ Cu^2+^ group showed 1.58-fold higher collagen I and 2.21-fold higher keratin 10 levels compared to the negative control, similar to the positive control (**Figure 7B**). Although the w/o Cu^2+^ group enhanced intensity compared with the negative control, the improvement was not statistically significant. In the skull region, immunostaining for osteogenic markers OPN and OCN was conducted (**Figure 7C**). Both scaffold-implanted groups improved expression of these markers relative to the negative control (**Figure 7D**). Notably, the w/ Cu^2+^ group showed significantly higher intensities in both OPN and OCN compared to the w/o Cu^2+^ group.

**Figure 7.**
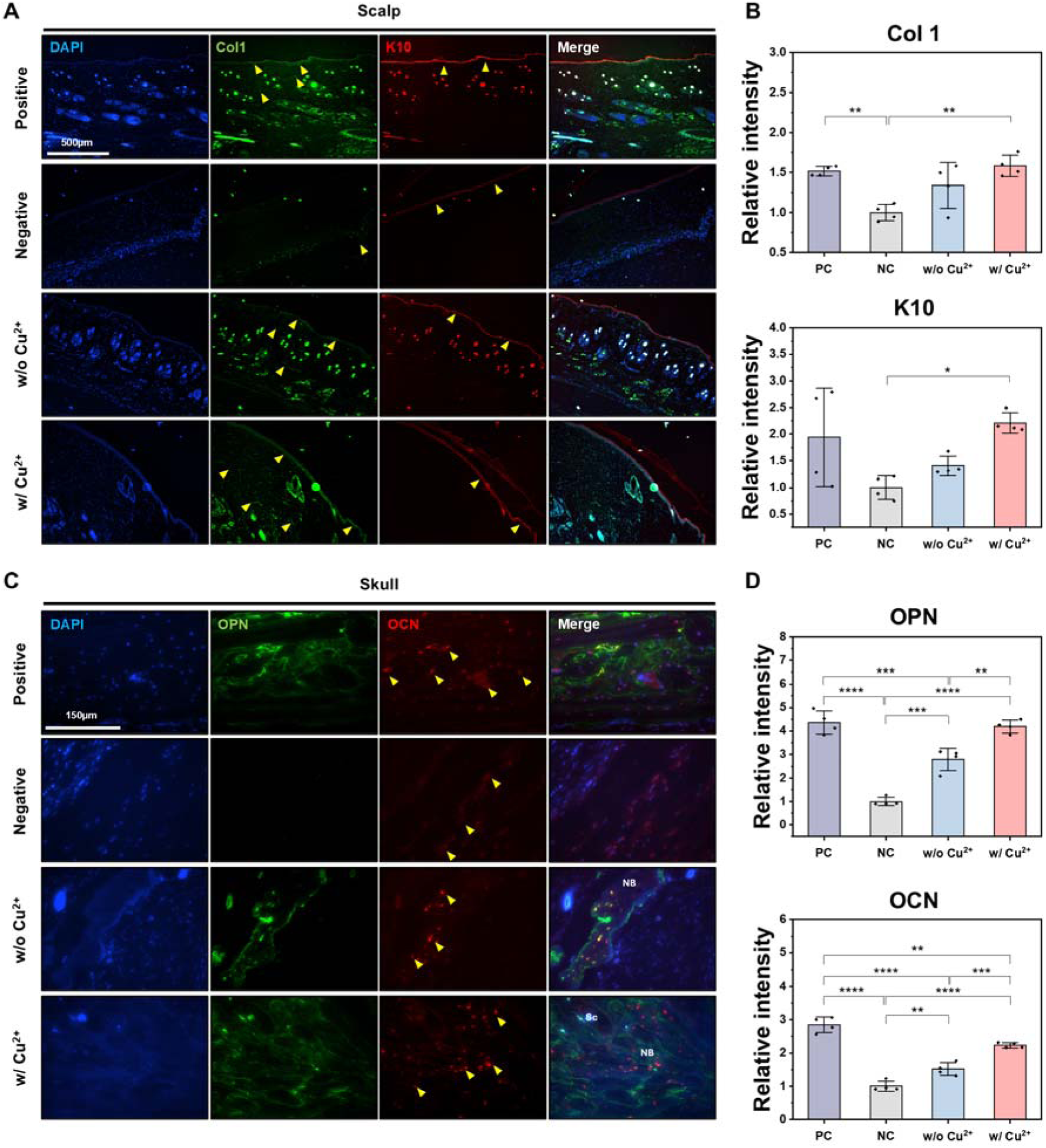
Skin tissue regeneration and osteogenic potential in the composite cranial defect model. **(A)** Representative immunofluorescence images of Collagen I (Col1) and Cytokeratin-10 (K10) in the scalp region at 12 weeks. Yellow arrow: positive staining. **(B)** Quantification of relative mean gray intensity for each marker in scalp tissue. **(C)** Representative immunofluorescence images of osteogenic markers OPN and OCN in the skull region at 12 weeks. **(D)** Quantification of relative mean gray intensity for each osteogenic marker. Yellow arrow: positive staining; Sc: scaffold; NB: newly formed bone.

Taken together, A5G81/Cu-PPCN hydrogels significantly enhanced blood vessel formation and promoted maturation of regenerated tissue (**Figure 7B**), despite no notable morphological difference being observed histologically in the scalp region (**Figure 6A**). Furthermore, BGCu-PPCN hydrogels not only induced more mature bone formation but also improved extracellular matrix deposition (OPN) and promoted mineralization (OCN) (**Figure 7D**).

## 3. Discussion

In clinical practice, treatment of composite cranial defects often requires multiple surgical stages. Surgeons typically prioritize scalp reconstruction, as inadequate soft tissue coverage can lead to severe complications such as implant exposure.^[32]^ Definitive cranial reconstruction is therefore often delayed until healthier soft tissue has formed. However, soft tissue coverage alone lacks sufficient structural support and can lead to certain negative sequelae, such as syndrome of the trephined.^[33]^ Synthetic materials used for bone reconstruction, such as PEEK and PMMA, continue to show high postoperative infection rates or insufficient support for bone ingrowth and vascularization.^[34]^ This challenge is amplified in open traumatic cranial defects, where the lack of a protective barrier increases susceptibility to bacterial invasion and delays healing.^[35, 36]^ Although numerous tissue engineering strategies have been proposed to reconstruct bone and soft tissues, many focus primarily on biological efficacy or single tissue type while overlooking practical clinical constraints. As a result, conventional scaffolds rarely meet the comprehensive biological and structural demands required for one-stage reconstruction of complex CMF composite defects. In particular, the incorporation of exogenous biological factors, despite demonstrating robust regenerative outcomes at the experimental scale, often limits clinical translation due to manufacturing complexity, cost, and regulatory burden.^[15]^ These considerations underscore the need for clinically appropriate material selection and strategy, and for an acellular, biocompatible, and integrated platform capable of regenerating composite tissues within a unified construct.

Our system was designed to address recurrent infection, inadequate vascularization, and the need for either complex multi-tissue vascularized free tissue transfer or multistaged surgeries in clinical management of composite defects. Importantly, all components are derived from citrate-based biomaterials (CBBs). CBBs possess inherent antioxidative and anti-inflammatory properties, and they have been used in FDA-cleared bioabsorbable implantable devices for use musculoskeletal reconstructive surgeries,^[37]^ highlighting their translational advantages. To mitigate potential Cu²□-related toxicity, we engineered controlled release by sequestering copper within BGCu or CMOF and integrating these nanoparticles into the CBB hydrogel (PPCN), enabling sustained local delivery.^[22, 38]^ Based on our *in vitro* release data, the *in vivo* formulation is estimated to release ∼1 µg of copper ions over 72 hours, corresponding to ∼15.7 µM if distributed within ∼1 mL of surrounding tissue/fluid. This level is within reported total copper ion concentrations in plasma (∼15 µM) and cerebrospinal fluid (∼50 µM). Although higher than brain extracellular free copper ion estimates (∼0.2–1.7 µM),^[39]^ *in vivo* copper is predominantly protein-bound, reducing the bioavailable free Cu²□ fraction.^[40, 41]^ Consistent with this finding, at 12-week, our CBB-based system showed no significant pro-inflammatory responses (**Figure S10**). Notably, only the copper ion-eluting groups induced significant M2 macrophage polarization, indicating an immunoregulatory shift consistent with a pro-healing environment.^[42, 43]^ This subsequent reduction in the immune response may be attributed to the antioxidant properties of CBBs,^[37]^ and citrate accumulation in mitochondria exerting indirect anti-inflammatory effect and positive bioenergetic effects.^[5, 44]^ The sustained low-dose Cu²□ release may also contribute to macrophage polarization toward an M2 phenotype by upregulating reparative signals.^[45]^

The distinct differences observed between BG and BGCu groups *in vitro* and *in vivo* highlight the contribution of copper ions to bone regeneration. Although BG coated mPOC-60HA scaffold showed higher osteogenic marker expression, BG coated scaffold resulted in incomplete mineralization and limited osteoid formation *in vivo* model. In contrast, BGCu markedly enhanced osteoid maturation and mineralization, leading to bone regeneration comparable to autologous grafts. This result was supported by earlier expression of late-stage osteogenic markers (OPN and OCN) *in vitro,* along with significantly higher expression levels at 12 weeks *in vivo*. Recent studies suggest that the copper ion facilitates osteoblast mitophagy and mitochondrial dynamics, which in turn enhance amorphous calcium phosphate release.^[46]^ Given that amorphous calcium phosphate is a key precursor for biomineralization, it aligns with the improved bone formation and higher mean HU values observed in w/ Cu^2+^ compared with the w/o Cu^2+^ treated groups. Successful peripheral bone bridging and defect coverage (e.g., defect size reduction) may be attributed to copper-mediated vessel-guided osteogenesis, whereby vascular ingrowth directs and sustains continuous bone formation across the defect.^[47]^ Another pivotal factor is that the antimicrobial efficacy of the BGCu suppressed bacterial colonization, mitigated infection-associated bone resorption and enhanced the stability of scaffold-tissue integration,^[48, 49]^ thereby preserving a permissive microenvironment for bone regeneration and mineralization. By contrast, the autograft showed only partial integration with the surrounding bone *in vivo*. This may be due to the absence of plate and screw fixation in our model, unlike standard clinical practice, as well as the intrinsic resorption tendency of autografts.^[50]^

A5G81 is a 12-amino-acid peptide derived from the α5-globular domain of laminin that promotes dermal tissue regeneration through α3β1 and α6β1 integrin-mediated cell interactions.^[21, 51]^ The absence of a significant difference between the A5G81-PPCN and PBS groups *in vitro* suggests the intrinsic wound-healing activity of A5G81 mediated through receptor interactions. While A5G81-PPCN facilitated early wound closure *in vivo* through direct cell-matrix interactions, the incorporation of CMOF provided additional antimicrobial properties and enhanced vascularization. Notably, the accelerated wound closure achieved with A5G81/Cu-PPCN was accompanied by histologically organized epidermal and dermal regeneration, with K10 and collagen I comparable to the autologous positive control and markedly higher than the untreated control, indicating improved healing in both rate and quality. These effects are likely attributable to Cu²□-induced angiogenesis, which improves oxygen and nutrient delivery, and may stem from copper ion-mediated regulation of epidermal cell migration and dermal-epidermal junction integrity.^[22, 52]^ In addition, Cu²□ serves as a co-factor for lysyl oxidase, indirectly promoting collagen crosslinking, which may contribute to matrix stability within the healing tissue.^[53]^ Collectively, these outcomes suggest synergistic interactions between biochemical peptide signaling (A5G81) and ion-mediated stimulation from Cu²□. Importantly, this synergy enabled A5G81/Cu-PPCN to support organized regeneration even within chronic composite defects, an environment in which insufficient perfusion typically impedes both re-epithelialization and matrix maturation.^[54, 55]^

Despite the efficacy of our scaffold demonstrated in this study for repairing complex composite defects, several limitations should be acknowledged. First, the evaluation was primarily conducted using a non-infected chronic defect model. Our *in vitro* assays and wound swab analyses demonstrated the antibacterial capacity of this system and its ability to suppress endogenous bacterial colonization. However, its therapeutic efficacy has not yet been directly evaluated in a bacteria-inoculated, high-burden infection model. Second, this study relied on a rat model. Given the inherent differences in bone metabolism rate, skin regeneration dynamics, anatomical scale, and immune response between rodents and humans,^[56, 57]^ future validation in large animal models is essential to assess the long-term stability of the system and regenerative performance under clinically relevant scales and biomechanical environments. Finally, the small sample size (n = 4 per group) may not have been sufficiently powered to detect small differences between groups. The copper ion-eluting treatment group showed marked improvements in bone formation and mineral density. Accordingly, interpretation was based on the magnitude of change relative to the negative and positive controls rather than p-values alone.

## 4. Conclusion

We demonstrate that a controlled copper ion-eluting, citrate-based acellular platform supports simultaneous regeneration of composite soft–hard tissue defects. Importantly, our system was evaluated in a chronic defect model,^[24]^ which more closely reflects clinically relevant traumatic and chronic composite defects. In this setting, it achieved composite tissue regeneration comparable to autologous tissue. We would predict that our regenerative strategy would reduce the donor site burden on a larger scale where autologous tissue options may be exhausted. This unified acellular copper ion-eluting system offers strong clinical potential for simplifying surgical procedures and achieving functional restoration of composite soft and hard tissue defects.

## Acknowledgements

This work was supported by the Northwestern University NUSeq Core Facility and made use of the IMSERC NMR and Physical Characterization facility at Northwestern University, which has received support from the Soft and Hybrid Nanotechnology Experimental (SHyNE) Resource (NSF ECCS-2025633), and Northwestern University. This work made use of the Keck-II facility and the EPIC facility of Northwestern University’s NUANCE Center, which has received support from the SHyNE Resource (NSF ECCS-2025633), the IIN, and Northwestern’s MRSEC program (NSF DMR-2308691). Imaging work was performed at the Northwestern University Center for Advanced Molecular Imaging (RRID:SCR_021192) generously supported by NCI CCSG P30 CA060553 awarded to the Robert H Lurie Comprehensive Cancer Center. The micro-CT imaging analysis was done at The University of Chicago Integrated Small Animal Imaging Research Resource (iSARIRR) Facility, which was supported by the National Center for Advancing Translational Sciences (NCATS) of the National Institutes of Health (NIH) through Grant Number 2UL1TR002389-06 that funds the Institute for Translational Medicine (ITM). This work was supported by NIH (1-R01 DE030480, RRR). This work was supported by the Querrey Simpson Institute for Regenerative Engineering at Northwestern University.

## Data Availability Statement

All data is available in the main text or the supplementary materials.

