## supplementary information for "Simultaneous regeneration of skin and bone in full-thickness cranial composite defects"

### Experimental Section

#### ***Synthesis and characterization of mPOC***

Methacrylated POC (mPOC) was developed via two-step synthesis process. First, POC pre-polymer was synthesized by adding equimolar amounts of citric acid and 1,8-octanediol in a round-bottom flask and melting the mixture at 165 °C. The temperature was then reduced to 140 °C, and the reaction was stirred for 50 min under a continuous nitrogen flow. The resulting pre-polymer was precipitated in Milli-Q (MQ) water and lyophilized to obtain POC. The lyophilized POC was dissolved in tetrahydrofuran at 60 °C, followed by the addition of imidazole and glycidyl methacrylate. The mixture was allowed to react under reflux for 6 h. After completion, the reaction mixture was concentrated using a rotary evaporator and precipitated in excess MQ water. The final mPOC product was collected and lyophilized for 4 days.

The chemical structure of mPOC was confirmed via $^{1}$H-NMR (X500) using DMSO-d6 as a solvent, and by FT-IR spectroscopy (Thermo Nicolet Nexus) in ATR mode.

#### ***Synthesis and characterization of PPCN and A5G81-PPCN***

PPCN was synthesized using a two-step procedure. First, the PPCac pre-polymer was prepared by melting citric acid, polyethylene glycol (PEG, Mw 400), and 1,3-diglycerolate diacylate at a molar ratio 5:9:1 at 140 °C under a nitrogen atmosphere. The reaction mixture was stirred for 40 min, after which the resulting PPCac pre-polymer was collected. In the second step, PPCac and N-isopropylacrylamide were mixed at a 1:1 molar ratio and dissolved in 1,4-dioxane at 65 °C. Azobis(isobutyronitrile) was added as a radical initiator, and the reaction proceeded overnight under nitrogen. After completion, the mixture was diluted with additional 1,4-dioxane, precipitated in diethyl ether, and vacuum-dried. The precipitate was neutralized to pH 7.4, dialyzed against MQ water at 4 °C, and lyophilized to obtain the final PPCN polymer.

For the synthesis of peptide-functionalized PPCN (A5G81–PPCN), PPCN was dissolved in MQ water at a concentration of 100 mg/mL, followed by addition of NHS (N-hydroxysuccinimide) and EDC (1-ethyl-3-(3-dimethylaminopropyl)carbodiimide hydrochloride) to activate the carboxyl groups. After 2 h of reaction at room temperature, BMPH (N-(β-maleimidopropionic acid) hydrazide) was added and allowed to react overnight at room temperature. The resulting BMPH-conjugated PPCN was adjusted to pH 7.4, dialyzed for 2 days at 4 °C, and lyophilized. For the final conjugation, PPCN, BMPH–PPCN and A5G81 peptide (0.01mmol) were mixed in PBS and reacted overnight at room temperature. The product was then dialyzed for 2 days at 4 °C and lyophilized prior to use.

The chemical structure of PPCN was confirmed via $^{1}$H-NMR (X500) using DMSO-d6 as a solvent, and by FT-IR spectroscopy (Thermo Nicolet Nexus) in ATR mode. Molecular weights of PPCN and the conjugation of A5G81–PPCN were evaluated via MALDI-TOF mass spectrometry, using α-cyano-4-hydroxycinnamic acid (CHCA) matrix dissolved in acetonitrile/water with 0.1% TFA.

#### ***Fabrication and characterization of BG and BGCu***

Bioactive glass was synthesized via a sol–gel process. Briefly, TEOS (Tetraethyl orthosilicate) (21.6 mL), MQ water (13.9 mL), and 2 M $HNO_3$ (2.8 mL) were dissolved in absolute ethanol (50 mL) under continuous stirring at room temperature for 1 h. Subsequently, TEP (Triethyl phosphate) (2.2 mL) was added and stirring continued for an additional 1 h. Then, calcium nitrate (14.04 g) was introduced into the solution and mixed until fully dissolved. For the synthesis of copper-doped bioactive glass, copper oxide (1.1 g) was added once the calcium nitrate had fully dissolved, followed by stirring for 1 h. After complete dissolution of all precursors, 1 M ammonia solution (10 mL) was gradually added to the acidic mixture under vigorous stirring until the solution transitioned to gelation. The resulting gel was incubated at 60 °C overnight to remove residual water and ethanol. The dried gel powder was subsequently heat-treated at 600 °C for 2 h to obtain the final bioactive glass and copper-doped bioactive glass.

The chemical structure of the product was confirmed using FT-IR (Thermo Nicolet Nexus) spectroscopy in both transmission mode with KBr pellets and ATR mode.

***Fabrication and characterization of CMOF particles***

Copper-based metal–organic framework (CMOF) particles were synthesized as follows. Copper acetate monohydrate (0.75 mmol) was dissolved in a solvent mixture of MQ water, ethanol, and DMF (1:1:1 v/v/v). Separately, $H_3BTC$ (trimesic acid) was dissolved in 6 mL of the same solvent mixture, and the two solutions were then combined. The reaction mixture was stirred continuously at room temperature overnight, with the vessel covered to minimize solvent evaporation. After completion, the mixture was centrifuged at 5000 rpm for 10 min to collect the blue precipitate. The supernatant was discarded, and the precipitate was washed sequentially with ethanol (×3) and MQ water (×3). The crude product was dried at 80 °C for 16 h, then stored in a desiccator to prevent moisture uptake.

The chemical structure of the synthesized CMOF particles was verified using FT-IR (Thermo Nicolet Nexus) spectroscopy in both transmission mode with KBr pellets and ATR mode. The crystalline phase of the material was determined via X-ray diffraction (XRD) using a Cu rotating-anode X-ray source. The morphology and particle size were evaluated using scanning electron microscopy (SEM) (SU-8030, Hitachi, Japan) after Au/Pt sputter coating.

***Preparation of composite PPCN hydrogel***

Each composite PPCN hydrogel was fabricated by dissolving PPCN precursor (100 mg/mL) in PBS, followed by incorporation of the respective components. For BG–PPCN and BGCu–PPCN composites, bioactive glass powder was mixed into the PPCN solution at a final concentration of 10 wt%. For A5G81/Cu–PPCN, CMOF particles were added to achieve a final concentration of 1 mM.

The lower critical solution temperature (LCST) of each hydrogel was evaluated using a discovery hybrid rheometer (TA Instruments, New Castle, DE). A total of 280 μL of precursor hydrogel solution was placed between parallel plates with a 1 mm gap. The storage modulus ($G'$) and loss modulus ($G''$) were recorded under oscillatory temperature ramp mode at a rate of 1 °C/min, within the range of 15−40 °C. All measurements were conducted at a constant frequency of 1 Hz and 1% strain. The LCST was defined as the intersection point of the G′ and G″ curves.

The release of calcium and copper ions from each composite hydrogel was quantified using inductively coupled plasma mass spectrometry (ICP-MS). The elemental composition of the composite hydrogels was analyzed by energy-dispersive X-ray spectroscopy (EDS) after Au/Pt sputter coating. The structural morphology was visualized using scanning electron microscopy (SEM) (SU-8030, Hitachi, Japan) after Au/Pt sputter coating.

***Fabrication of multilayered 3D-printed composite scaffold***

To prepare the composite inks for 3D printing, the mPOC polymer was dissolved in absolute ethanol at a final concentration of 80 wt%. For the skin layer and the interfacial transition layer, mPOC was dissolved in ethanol and supplemented with Irgacure 819 (1wt%) as the photoinitiator. For the bone layer, mPOC was mixed with hydroxyapatite (HA) nanoparticles at a 40:60 polymer-to-HA mass ratio, resulting in a composite ink containing 60 wt% solids, and Irgacure 819 (1wt%) was also added to the bone-layer ink.

Composite scaffolds were fabricated using a micro-continuous liquid interface production (μCLIP) 3D printer. The scaffold models were designed using SolidWorks and exported as CAD files for printing. The skin-layer scaffold was designed with a diameter of 6mm, height of 1.5 mm, and strut thickness of 125 μm. The bone layer was designed with a diameter of 4 mm and height of 1 mm. An interfacial transition layer was implemented as a 6mm-diameter disk containing 20 μm pores to facilitate integration between the skin and bone layers. These CAD models were sliced into cross-sectional images via the use of a homemade MATLAB code with 5μm layer thickness. The resulting cross-sectional images could then be projected onto the resin bath via the use of a digital micromirror device (DMD) with a wavelength of 365nm.

For the fabrication of multilayered composite scaffolds, the skin and interfacial transition layers (mPOC-based) were printed first using resin bath #1 containing mPOC ink. Subsequently, the bone layer (mPOC–60HA) was printed by replacing the resin bath #2 with the mPOC–60HA composite ink. Following printing, the scaffolds were washed with ethanol to remove unreacted resin and residual photoinitiator. The scaffolds were then post-cured under UV light for 1 min. Finally, the printed multilayered scaffold was coated with composite PPCN hydrogel at 37 °C.

***Cytocompatibility test***

Human bone marrow–derived mesenchymal stem cells (hMSCs; PCS-500-012, ATCC) were used for all in vitro studies. Cells were cultured in growth medium (PCS-500-030, ATCC) supplemented with the growth kit (PCS-500-040, ATCC) and maintained at 37 °C in a humidified incubator with 5% $CO_2$. Prior to cell seeding, all scaffold samples were sterilized using an ethylene oxide sterilization system and subsequently preconditioned by immersion in growth medium for 24 h at 37 °C.

For biocompatibility assessment, hMSCs (passage 5) were seeded at a density of 20,000 cells per well in 48-well plates. Scaffolds were then transferred into each well using a Transwell insert system. For hydrogel-coated composite scaffolds, a total of 100 μL of composite hydrogel precursor solution (50 μL of BG- or BGCu-PPCN and 50 uL of A5G81-

or A5G81/Cu-PPCN) was applied onto the scaffold prior to culture. Cell viability was evaluated at determined time points using the alamarBlue assay and Live/Dead staining, following the manufacturer's instructions. Cells cultured directly on a 2D tissue culture plate were used as the normalization control.

***In vitro* immunofluorescence staining of osteogenic markers**

hMSCs with low passage number (≤4) were used to assess osteogenic differentiation. Prior to induction, cells were cultured in growth medium for 48 h, after which the medium was replaced with osteogenic differentiation medium (PCS-500-052, ATCC).

For immunofluorescence evaluation of osteogenic marker expression, 40,000 cells per well were seeded in 48-well plates, and hydrogel-coated composite mPOC-60HA scaffolds were introduced into each well using a Transwell insert system. Cells were cultured for 7 and 14 days, then fixed with 4% paraformaldehyde and immunostained for RUNX2 (SC-390351, Santa Cruz), OPN (SC-21742, Santa Cruz Biotechnology), OCN (23418-1-AP, Proteintech), and counterstained with DAPI.

Fluorescence imaging was performed using a Nikon Eclipse Ti2-E microscope equipped with NIS-Elements software. The relative mean gray intensity of each marker was quantified using ImageJ software and normalized to the signal from the mPOC-60HA scaffold-only condition at day 7.

***Scratch assay***

Human epidermal keratinocytes (HEKa; passages 1–2) were used for the scratch assay. Cells were seeded at a density of $5 \times 10^4$ cells per well in 24-well plates and incubated overnight to allow cell attachment and monolayer formation. A linear scratch was introduced using a sterile pipette tip, and cells were then incubated with PBS (control), A5G81–PPCN hydrogel, or A5G81/Cu–PPCN hydrogel. Digital images of the wounded area were captured after 36 h of incubation. The percentage of wound closure was calculated using the following equation:

Wound closure (%) = $(A_{initial} - A_{final}) / A_{initial} \times 100\%$

Where $A_{initial}$ is the scratch area immediately after scratching and $A_{final}$ is the scratch area after 36 h.

***Tubulogenic assay***

Human umbilical vein endothelial cells (HUVECs; passages ≤5) were seeded in 24-well plates at a density of $5 \times 10^4$ cells per well and cultured overnight to allow cell attachment. Cells were then treated with PBS (control), BG–PPCN, BGCu–PPCN, A5G81–PPCN, or A5G81/Cu–PPCN hydrogels for 24 h. Matrigel (Corning, CLS354234) was diluted 1:1 with vascular basal medium, and 60 µL of the mixture was added to each well of a 96-well plate, followed by incubation at 37 °C for 30 min to allow gelation. After treatment, cells were detached, resuspended in basal medium, and seeded onto the Matrigel at a density of $2.5 \times 10^4$ cells per well, followed by incubation for 4 h at 37 °C. Formation of enclosed capillary-like tube networks was digitally imaged, and angiogenic parameters were quantified using the Angiogenesis Analyzer plugin in ImageJ.

***Antibacterial assay***

Staphylococcus aureus (S. aureus) was cultured in LB broth at 37 °C and 200 rpm overnight. The bacteria were harvested, washed, and resuspended in PBS at a concentration of 1 × $10^5$ CFU/mL. A total of 100 µL of composite hydrogel (prepared in PBS) was added to each well of a 96-well plate, followed by addition of the bacterial suspension. The plate was then incubated at 37 °C for 24 h. After incubation, 100 µL of the bacterial suspension was collected from each well and serially diluted in sterile PBS. A volume of 20 µL from each diluted sample was spread onto LB agar plates and incubated at 37 °C overnight. Digital images of the agar plates were acquired, and the colony-forming units (CFU) were quantified using ImageJ software. All experiments were conducted in triplicate (n = 3). The bacterial killing ratio was calculated as follows:

Killing ratio = (CFU of negative control – CFU of hydrogel group) / CFU of negative control x 100%

***Surgical procedures***

The protocols of this study were approved by the Institutional Animal Care and Use Committee (IACUC) of University of Chicago (ACUP #71445). Bone and soft tissue defects in rat models (8-week-old Sprague Dawley rats, weight range 200-350g; Envigo, Indianapolis, IN) were carried out following the approved guidelines.

Anesthesia was induced in rats through isoflurane inhalation, complemented by intraperitoneal injection of ketamine (50–75 mg/kg) and xylazine (7–10 mg/kg). After prone positioning, the rat was prepared by scalp depilation, sterilization with iodine tincture, and application of a sterile drape. Using a biopsy punch, two 6 mm circular incision was made above the bilateral parietal cranium, and the pericranium was removed with a neuro spatula. A cranial defect was then carefully drilled in the parietal bone with a 4 mm trephine under continuous normal saline irrigation to prevent tissue thermal injury (Dremel® USA, Robert Bosch Tool Corp). The bone disc was removed, and hemostasis was meticulously achieved. A wound obturator was implanted in each wound to counteract the rapid initial self-healing of the scalp post-trauma1. Autologous bone discs were preserved in saline-soaked gauze at 4 °C. Three weeks after the initial surgery, chronic wound models were established. The wound obturators were removed, and the surrounding soft tissues were carefully debrided. The left-side defects were designated as either the negative control (no treatment) or the positive control (reimplantation of autologous bone followed by skin suture), whereas the right-side defects were assigned to the experimental scaffold groups, with or without $Cu^{2+}$ incorporation.

***Micro-CT imaging and analysis***

Live rats were subjected to micro-CT scans at weeks 0 (three days post implantation operation), 3, and 12, using the X-CUBE Preclinical CT Imager (Molecubes NV, Belgium) at The University of Chicago Integrated Small Animal Imaging Research Resource (iSAIRR) facilities. Spiral high-resolution CT acquisitions were performed with an x-ray source of 50 kVp and 200 µA. Volumetric CT images were reconstructed in a 350 × 350 × 840 format with voxel dimensions of 200 µm3. A few volumetric CT images were also reconstructed in a 700 × 700 × 374 format with voxel dimensions of 100 µm3 to evaluate the performance of bone healing. Reconstructions were performed using 3-D Slicer software (Version 5.4.0) as previously described 2-8. Defect areas were quantified using Image J (Version 1.53k).

Relative Defect Area Ratio (RDAR), relative bone volume and mean HU was calculated to quantify the healing rate. RDAR= (Area at each timepoint/Area at week 0) x 100%.

***In vivo wound healing assessment***

Wound healing was assessed and documented daily until complete epithelial closure. For the negative control group (left side), the Tegaderm dressing was removed once scab formation was observed. For the scaffold-treated group (right side), the wound surface was refreshed and covered with freshly prepared A5G81-PPCN or A5G81/Cu-PPCN hydrogels every two days. Wound closure was recorded photographically, and the wound area was quantitatively analyzed using ImageJ (Version 1.53k).

On postoperative day 5, sterile wound cotton swabs were used to collect exudate samples from the wound bed. Each swab was rinsed three times in 1 mL of LB broth, and 100 µL of the suspension was plated onto antibiotic-free LB agar plates. After incubation at 37 °C for 24 hours, bacterial growth was documented by photography, and colony numbers were quantified.

***Histological analysis***

At 12 weeks post-implantation, all rats were euthanized, and tissue samples were harvested, fixed and decalcified using Cal-Ex II fixative/decalcifier solution (Fisher Chemical). Decalcification was continued at 4°C until the cranial bone could be easily penetrated using a micro-needle, after which samples were rinsed twice with PBS. The tissues were then dehydrated through a graded ethanol series, with each step lasting 90 min, followed by clearing in xylene and paraffin embedding.

For histological evaluation, paraffin-embedded tissue blocks were sectioned at 5 µm thickness. Sections were stained with hematoxylin and eosin (H&E) and Masson's trichrome (Masson Trichrome Kit 87019, Epredia) to assess biocompatibility and tissue formation. Stained sections were examined using a bright-field microscope (Eclipse Ti2-E, Nikon).

***Immunofluorescence tissue staining***

For IF staining, paraffin-embedded tissue sections were subjected to heat-induced epitope retrieval, followed by blocking with 1% bovine serum albumin (BSA) in PBS at room temperature for 30 min. After blocking, sections were incubated overnight at 4 °C with the following primary antibodies: CD86 (14-0862-82, Invitrogen), CD163 (ab182422, Abcam), CD31 (ab182981, Abcam), α-SMA (α-Smooth Muscle Actin; ab7817, Abcam), Collagen I (Col1) (PA5-29569, Invitrogen), Cytokeratin 10 (K10) (MA5-13705, Invitrogen), RUNX2 (SC-390351, Santa Cruz), OPN (SC-21742, Santa Cruz Biotechnology), OCN (23418-1-AP, Proteintech). After primary antibody incubation, sections were washed and incubated with the appropriate Alexa Fluor–conjugated secondary antibodies for 30 min, followed by DAPI counterstaining to visualize nuclei.

Imaging was performed using a Nikon Eclipse Ti2-E microscope equipped with NIS-Elements software. Relative mean gray intensity was quantified using ImageJ, with the negative control group used for normalization for Col1, K10, OPN, OCN, CD86, and CD163.

Blood vessel density was quantified as the number of vessels per $mm^2$ within each tissue region. Macrophage polarization was evaluated using the M2/M1 ratio, normalized to the negative control.

***Statistical analysis***

Mean and standard deviation values were reported in this study. Statistical significance of differences between two conditions was assessed using Student's t-test. Statistical significance of difference between multiple conditions was evaluated by one-way ANOVA followed by Tukey's post hoc test. p-values below 0.05 were considered statistically significant and are reported here.

### Supplementary Figures

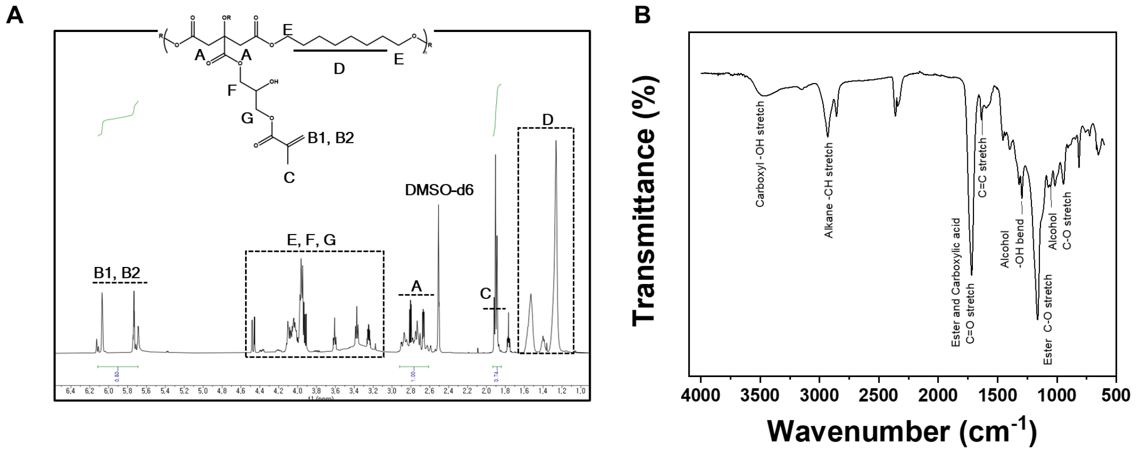


**Figure S1. Representative $^1$H-NMR spectra and FT-IR spectra of mPOC polymer. (A)** Characteristic peaks corresponding to POC, and methacrylate group are assigned in $^1$H-NMR spectra. **(B)** In FT-IR spectra, various functional groups within the mPOC polymer are assigned.

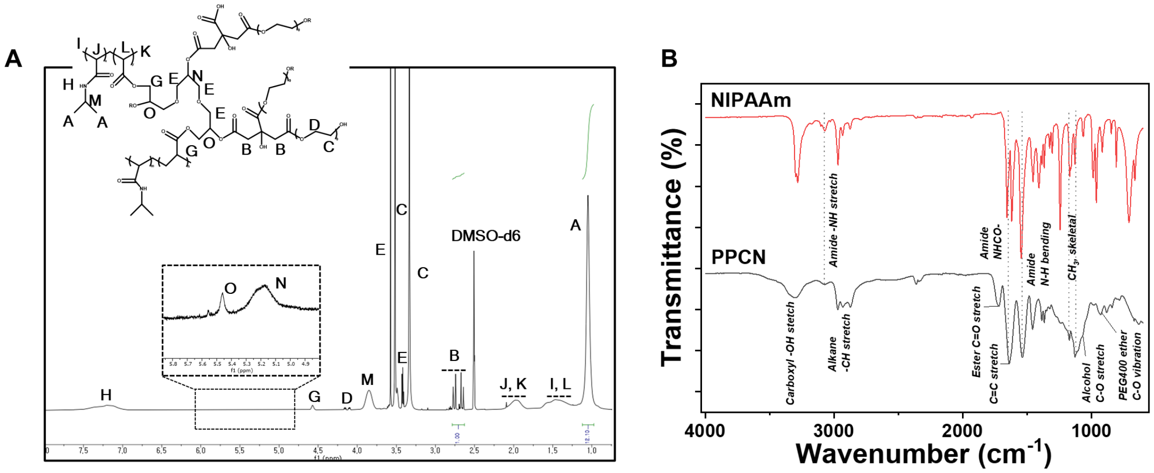


**Figure S2. Representative ¹H-NMR spectra and FT-IR spectra of PPCN polymer. (A)** The ¹H-NMR spectra identify and assign characteristic peaks corresponding to PNIPAAm, PEG, and citric acid are assigned. **(B)** Identification and characterization of different functional groups within PNIPAAm and PPCN polymers by FT-IR analysis.

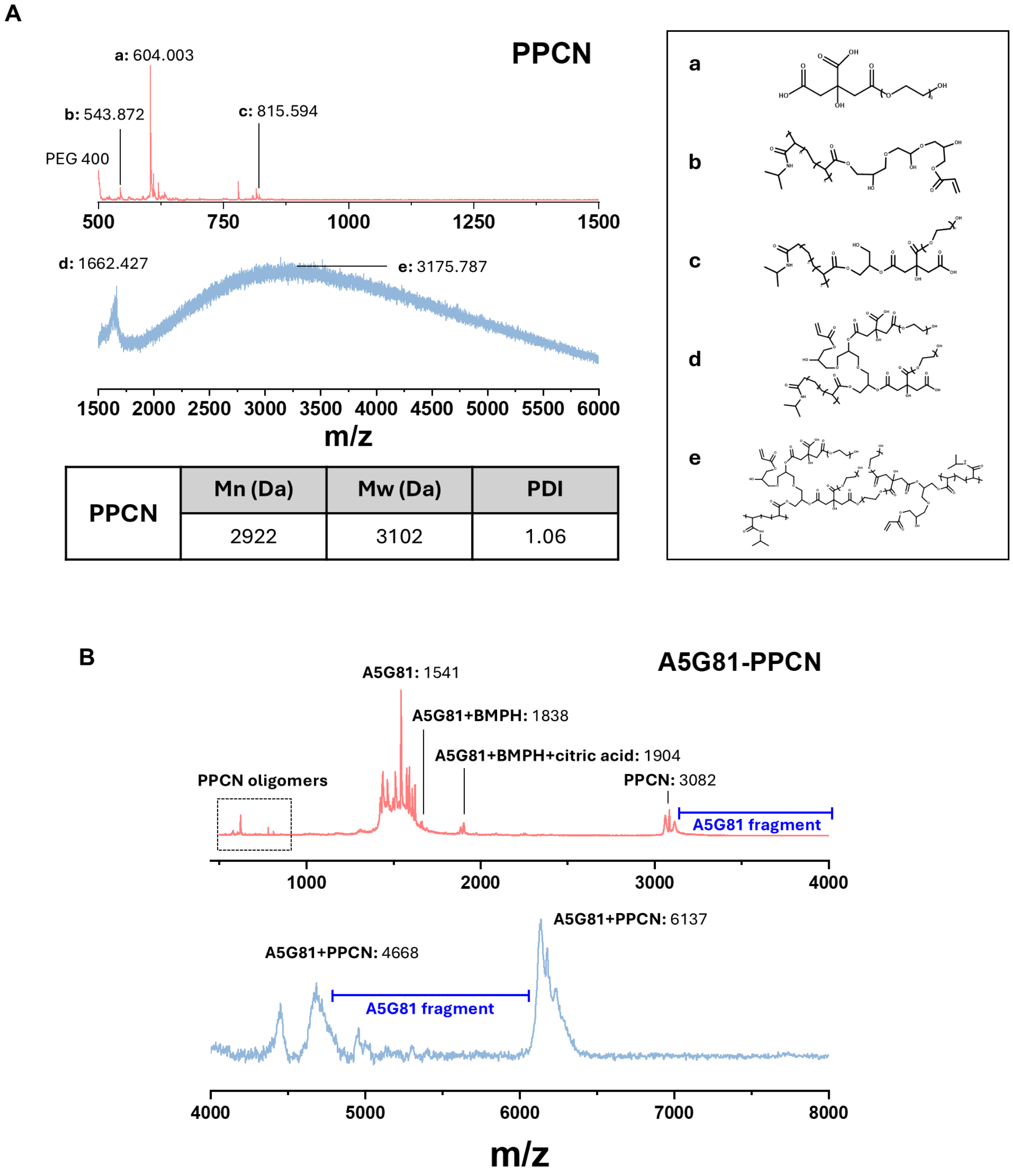


| | Mn (Da) | Mw (Da) | PDI |
|---|---|---|---|
| PPCN | 2922 | 3102 | 1.06 |

**Figure S3. Representative MALDI-TOF spectra of PPCN and A5G81-conjugated PPCN. (A)** MALDI-TOF spectrum of PPCN showing characteristic oligomer peaks. **(B)** MALDI-TOF spectrum of A5G81-PPCN showing characteristic oligomer peaks corresponding to A5G81-conjugated fragments.

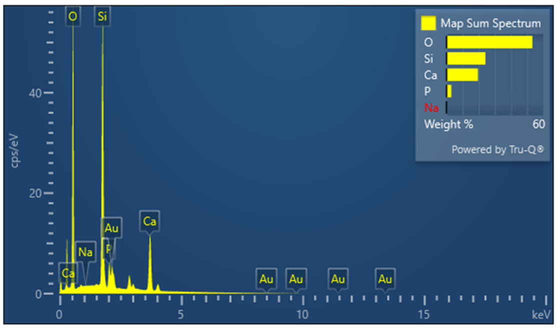


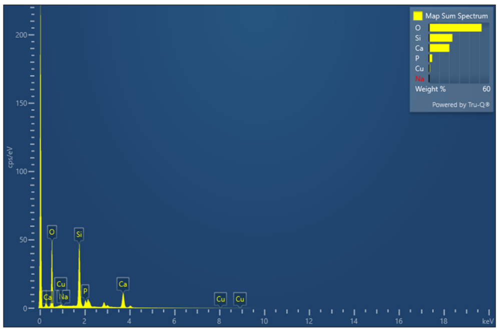


| **Bioactive glass** | | |
|---|---|---|
| **Element** | Wt% | Atomic % |
| **O** | 53.28 | 69.73 |
| **Si** | **24.17** | **18.02** |
| **Ca** | **19.52** | **10.20** |
| **P** | **3.00** | **2.03** |
| **Total** | 100.00 | 100.00 |

| **Copper doped-Bioactive glass** | | |
|---|---|---|
| **Element** | Wt% | Atomic % |
| **O** | 52.39 | 69.33 |
| **Si** | **23.30** | **17.56** |
| **Ca** | **20.22** | **10.68** |
| **P** | **3.05** | **2.09** |
| **Cu** | **1.03** | **0.34** |
| **Total** | 100.00 | 100.00 |

**Figure S4. Energy-dispersive X-ray spectroscopy (EDS) analysis of elemental composition in bioactive glass and copper-doped bioactive.** Each data shows elemental composition of bioactive glass and copper-doped bioactive glass powders.

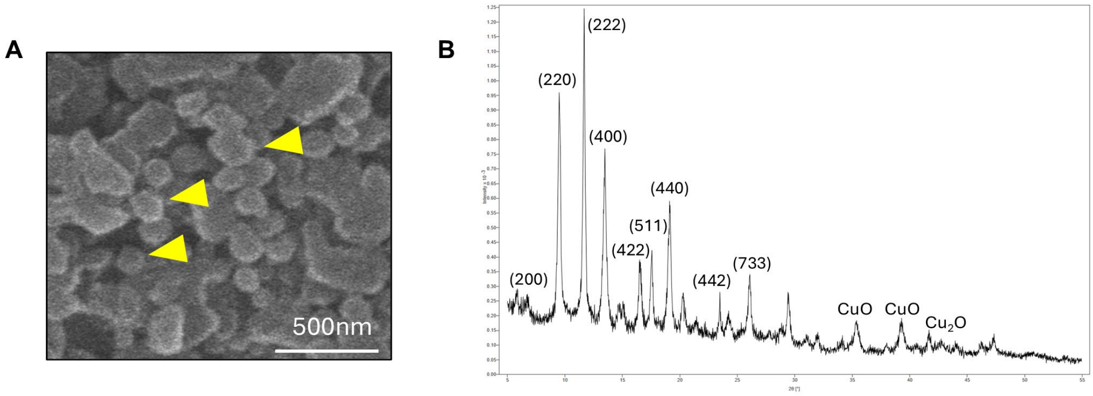


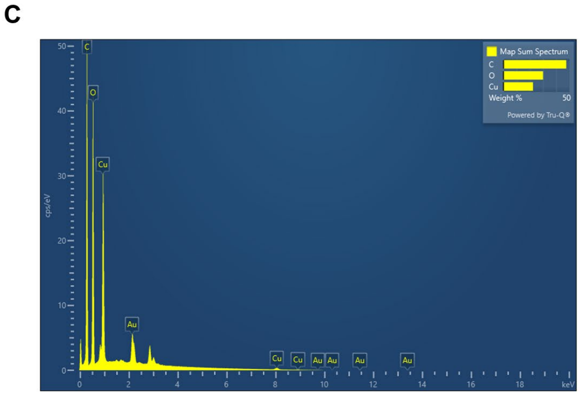


| Copper-MOF particles | | |
|---|---|---|
| **Element** | Wt% | Atomic % |
| **C** | 47.56 | 63.98 |
| **O** | 30.02 | 30.32 |
| **Cu** | **22.42** | **5.70** |
| **Total** | 100.00 | 100.00 |

**Figure S5. Characterization of Cu-MOF nanoparticles. (A)** SEM image of Cu-MOF nanoparticles. **(B)** X-ray diffraction (XRD) pattern of Cu-MOF. **(C)** Elemental composition of bioactive glass and copper-doped bioactive glass powders.

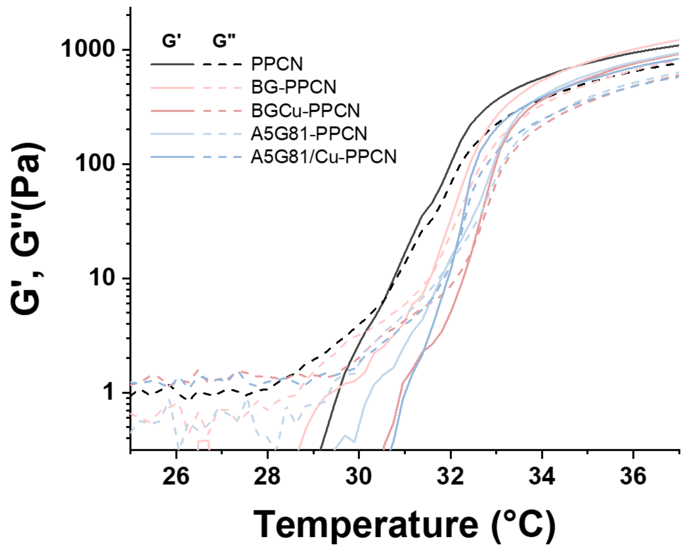


| PPCN | BG-PPCN | BGCu-PPCN | A5G81-PPCN | A5G81/Cu-PPCN |
|---|---|---|---|---|
| 30.54°C | 31.6°C | 32.44°C | 32.02°C | 32.23°C |

332 **Figure S6. Rheological analysis of PPCN and composite PPCN hydrogels.** Gelation
333 temperature (LCST) determined by the crossover point of G′ and G″ for each hydrogel.
334

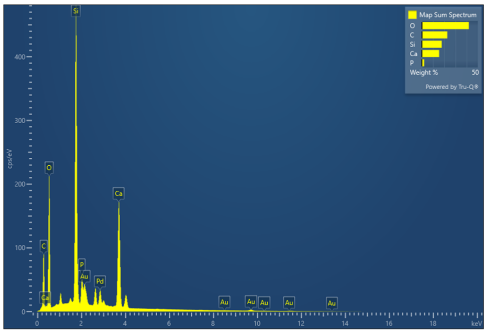


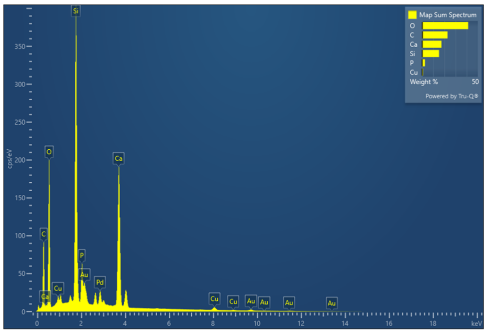


| Bioactive glass + PPCN | | |
|---|---|---|
| **Element** | Wt% | Atomic % |
| **C** | 22.75 | 33.81 |
| **O** | 41.94 | 46.8 |
| **Si** | **17.76** | **11.29** |
| **Ca** | **15.43** | **6.87** |
| **P** | **2.12** | **1.22** |
| **Total** | 100.00 | 100.00 |

| Copper doped-bioactive glass + PPCN | | |
|---|---|---|
| **Element** | Wt% | Atomic % |
| **C** | 20.3 | 30.77 |
| **O** | 43.12 | 49.08 |
| **Si** | **17.03** | **11.04** |
| **Ca** | **15.67** | **7.12** |
| **P** | **2.87** | **1.69** |
| **Cu** | **1.01** | **0.29** |
| **Total** | 100.00 | 100.00 |

**Figure S7. Energy-dispersive X-ray spectroscopy (EDS) analysis of elemental composition in bioactive glass and copper-doped bioactive composite hydrogels.** Each data shows elemental composition of the corresponding composite hydrogels.

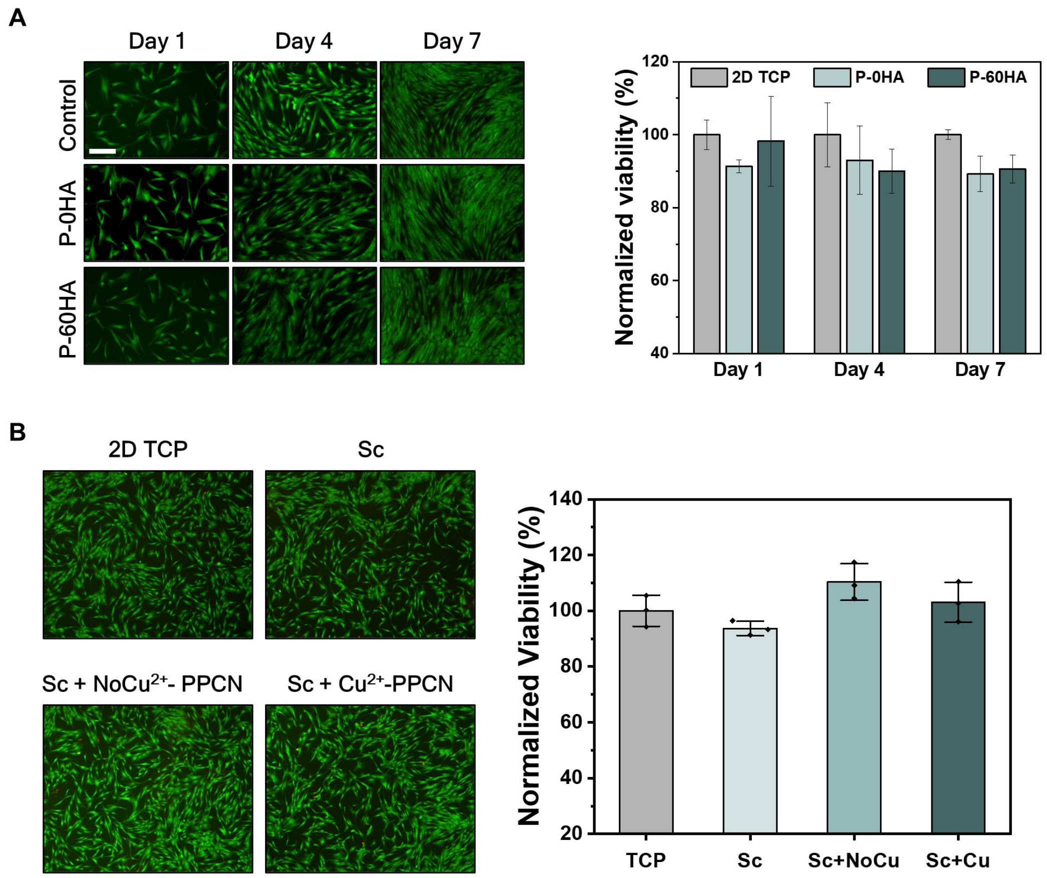


**Figure S8. In vitro assessment of hMSC viability on each scaffold system. (A)** Live/dead staining images and AlamarBlue assay (normalized to the TCP control) for 3D-printed scaffolds at days 1, 4, and 7. Scale bar: 200 µm. P-0HA: mPOC scaffold; P-60HA: mPOC-60HA scaffold. **(B)** Live/dead staining images and AlamarBlue assay (normalized to the TCP control) for hydrogel-coated composite scaffolds at days 1, 4, and 7. TCP: 2D tissue culture plate; Sc: Multilayered scaffold (P-0HA + P-60HA); Sc+NoCu$^{2+}$-PPCN: Multilayered scaffold coated with BG-PPCN and A5G81-PPCN; Sc+Cu$^{2+}$-PPCN: Multilayered scaffold coated with BGCu-PPCN and A5G81/Cu-PPCN.

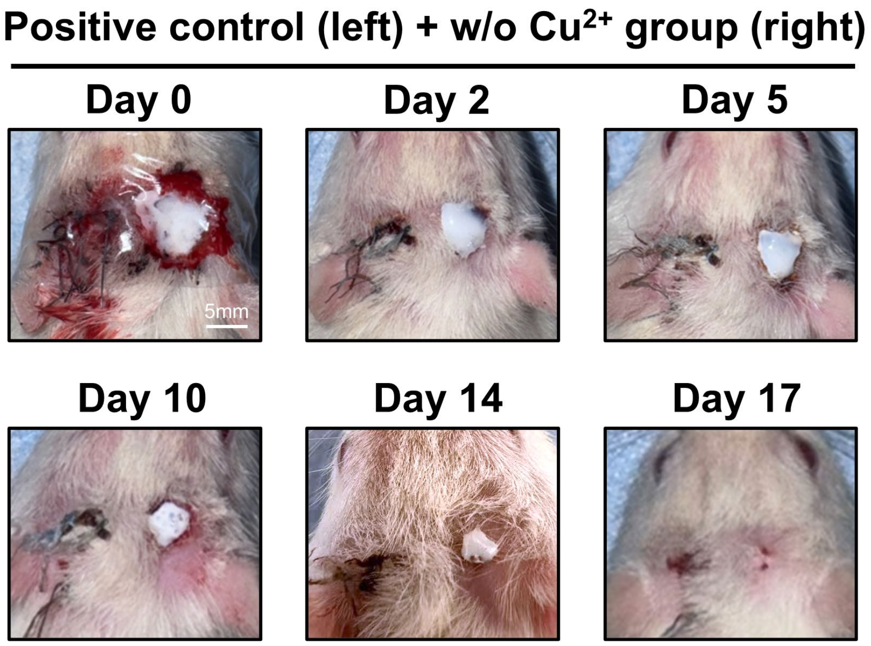


**Figure S9. Representative images of wound healing progression over time.** The positive control and non-copper–treated groups within the same biological model demonstrate healing progression until full closure.

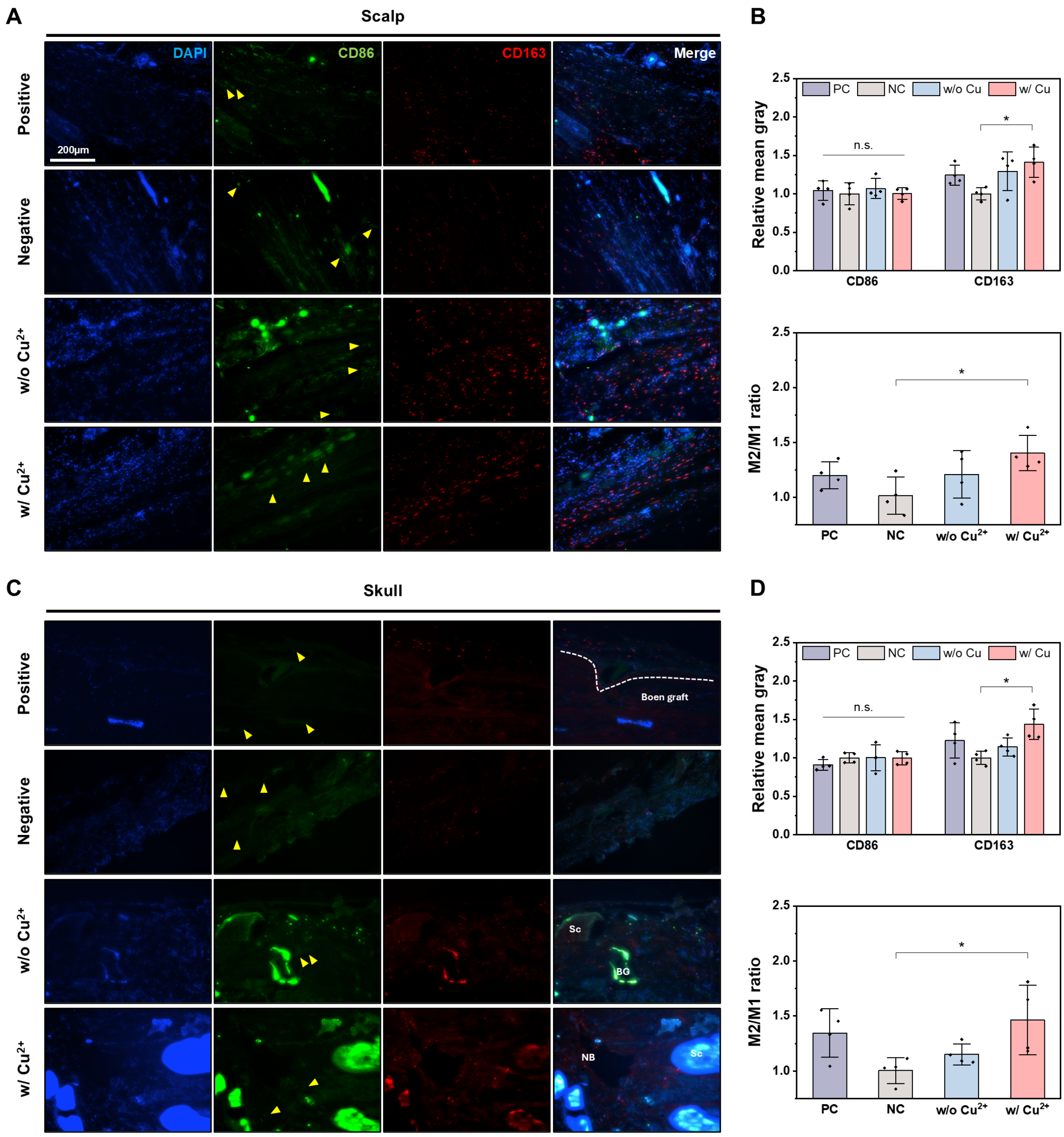


**Figure S10. Immunofluorescence staining of CD86 (M1) and CD163 (M2) markers in each tissue region. (A)** Representative immunofluorescence images of each group in the scalp region. Yellow arrows: CD86-positive cells. **(B)** Quantification of relative mean gray values for each marker and the corresponding M2/M1 ratio in the scalp region. **(C)** Representative immunofluorescence images of each group in the skull region. **(D)** Quantification of relative mean gray values for each marker and the corresponding M2/M1 ratio in the skull region.
